# Genome-wide association study reveals multiple loci for nociception and opioid consumption behaviors associated with heroin vulnerability in outbred rats

**DOI:** 10.1101/2024.02.27.582340

**Authors:** Brittany N. Kuhn, Nazzareno Cannella, Apurva S. Chitre, Khai-Minh H. Nguyen, Katarina Cohen, Denghui Chen, Beverly Peng, Kendra S. Ziegler, Bonnie Lin, Benjamin B. Johnson, Thiago Missfeldt Sanches, Ayteria D. Crow, Veronica Lunerti, Arkobrato Gupta, Eric Dereschewitz, Laura Soverchia, Jordan L. Hopkins, Analyse T. Roberts, Massimo Ubaldi, Sarah Abdulmalek, Analia Kinen, Gary Hardiman, Dongjun Chung, Oksana Polesskaya, Leah C. Solberg Woods, Roberto Ciccocioppo, Peter W. Kalivas, Abraham A. Palmer

## Abstract

The increased prevalence of opioid use disorder (OUD) makes it imperative to disentangle the biological mechanisms contributing to individual differences in OUD vulnerability. OUD shows strong heritability, however genetic variants contributing to vulnerability remain poorly defined. We performed a genome-wide association study using over 850 male and female heterogeneous stock (HS) rats to identify genes underlying behaviors associated with OUD such as nociception, as well as heroin-taking, extinction and seeking behaviors. By using an animal model of OUD, we were able to identify genetic variants associated with distinct OUD behaviors while maintaining a uniform environment, an experimental design not easily achieved in humans. Furthermore, we used a novel non-linear network-based clustering approach to characterize rats based on OUD vulnerability to assess genetic variants associated with OUD susceptibility. Our findings confirm the heritability of several OUD-like behaviors, including OUD susceptibility. Additionally, several genetic variants associated with nociceptive threshold prior to heroin experience, heroin consumption, escalation of intake, and motivation to obtain heroin were identified. *Tom1*, a microglial component, was implicated for nociception. Several genes involved in dopaminergic signaling, neuroplasticity and substance use disorders, including *Brwd1*, *Pcp4, Phb1l2* and *Mmp15* were implicated for the heroin traits. Additionally, an OUD vulnerable phenotype was associated with genetic variants for consumption and break point, suggesting a specific genetic contribution for OUD-like traits contributing to vulnerability. Together, these findings identify novel genetic markers related to the susceptibility to OUD-relevant behaviors in HS rats.

## Introduction

Opioid use disorder (OUD) is characterized by compulsive opioid use and relapse propensity, despite efforts to maintain abstinence. OUD diagnosis has increased substantially in the past decade (1), affecting approximately 16 million people per year worldwide (2). The rise in diagnosis is not demographically constrained (1), but is prevalent in individuals across socio-economic class, age and sex (3), increasing the urgency to better understand the biological mechanisms contributing to OUD vulnerability. In twin studies, OUD shows high heritability (∼50%) (4, 5), indicating that specific genetic variants contribute to biological basis of OUD susceptibility.

Genome-wide association studies (GWAS) are a useful tool for identifying the genetic variants that influence heritable traits contributing to OUD. GWAS of OUD, though limited, have already yielded valuable insights into genetic contributions to diagnostic vulnerability (6–13). However, human GWAS of OUD are limited to observational data and are thus confounded by numerous poorly defined environmental factors. Understandably, due to a lack of empirical data from study participants, human GWAS have largely focused on OUD diagnosis, which is comprised of poorly quantified amounts of drug-taking, extinction and seeking behaviors. Using rodent models of OUD, we can quantify specific traits (e.g., consumption, escalation, reinstatement) associated with addiction-related behaviors to directly interrogate the relationship between these metrics and genetic variants. By doing so, we can discover novel genetic variants that may contribute to the different phases of OUD, which can then be compared to data from human studies. Crucially, such genes may also provide targets for the development of novel OUD treatments.

In the current study, we assessed several measures of heroin-taking, extinction and seeking behaviors in outbred male and female heterogeneous stock (HS) rats at two different testing locations. Additionally, anxiety- and stress-like behaviors, as well as nociceptive threshold, were assessed prior to and following heroin experience. HS rats are a diverse outbred population that was created by interbreeding 8 inbred strains at NIH in 1984 to capture genetic and behavioral heterogeneity reflective of the human population (14–16). Compared to inbred and commercially available outbred rat lines, HS rats exhibit comparatively greater diversity (16) and have been used in several studies assessing addiction-related behaviors (17–24), including those focusing on genetic analyses (25–27). In addition, HS rats are supported by complementary expression quantitative trait loci datasets (28), furthering the translational utility of our findings. In this study, we found genetic variants specific to different OUD behavioral traits, many of which had known involvement in neurobiological processes associated with substance use disorder (SUD). Additionally, we employed a novel non-linear network-based clustering approach to identify subpopulations of OUD vulnerable and resilient rats (23, 24) and thereby identify genetic variants associated with OUD susceptibility.

## Methods

Behavioral testing occurred at two geographically distinct locations: Medical University of South Carolina (MUSC, USA) and the University of Camerino (UCAM, Italy). Two testing locations rendered the advantage of accounting for environmental differences on behavior, akin to the human experience, while maintaining standardized experimental procedures and equipment used between the sites. All experimental procedures were approved by the MUSC Institutional Animal Care and Use Committee and the Italian Ministry of Health (UCAM). All testing procedures were compliant with the National Institute of Health Guide for the Care and Use of Laboratory Animals and the Assessment and Accreditation of Laboratory Animals Care, as well as the European Community Council Directive for Care and Use of Laboratory Animals.

### Subjects

Male and female HS rats (NMcwiWFsm #13673907, RRID:RGD13673907) bred at Wake Forest University were shipped to MUSC and UCAM in batches of 40 (20 females and 20 males/site) at 5 weeks of age. Each site had an equal representation of littermates thereby creating genetically comparable testing cohorts. Whenever possible, each shipment contained only one male and one female per litter, to reduce genetic relationships, which can reduce power in GWAS. Upon arrival, rats were left undisturbed for 3 weeks in a climate-controlled vivarium room with a standard 12-hr light/dark cycle prior to testing. Rats were pair-housed and received ad libitum access to food and water throughout the duration of testing. Behavioral tests for stress, anxiety and nociception occurred during the light cycle, whereas all heroin-associated testing occurred during the dark phase. A total of 1,160 rats underwent testing from February of 2019 through December of 2022, with the following animals excluded from all analyses: 147 for behavioral control studies (e.g., did not undergo heroin self-administration procedures), 107 due to death (post-surgical illness, n=84; surgery, n=23); 29 for behavioral data collection technical issues, and 3 for spleen identification issues. Final behavioral and genetic analyses were conducted on 874 rats (MUSC, n=479; UCAM, n=395).

### Drugs

Heroin hydrochloride was supplied to both sites by the National Institute on Drug Abuse (Bethesda, MD) and dissolved in 0.9% sterile saline.

### Behavioral testing

The experimental timeline is shown in Figure 1a. Experimental testing occurred as previously described (17, 23, 24). Extensive behavioral analyses for these animals, including sex and site differences, have been conducted and reported in prior publications (17, 23, 24), and we refer the reader to these manuscripts for further edification. The focus of this manuscript is on identifying the genetic variants that contribute to individual differences in these behaviors. A brief description of behavioral testing procedures is provided below with greater detail in the Supplemental Information.

**Figure 1.**
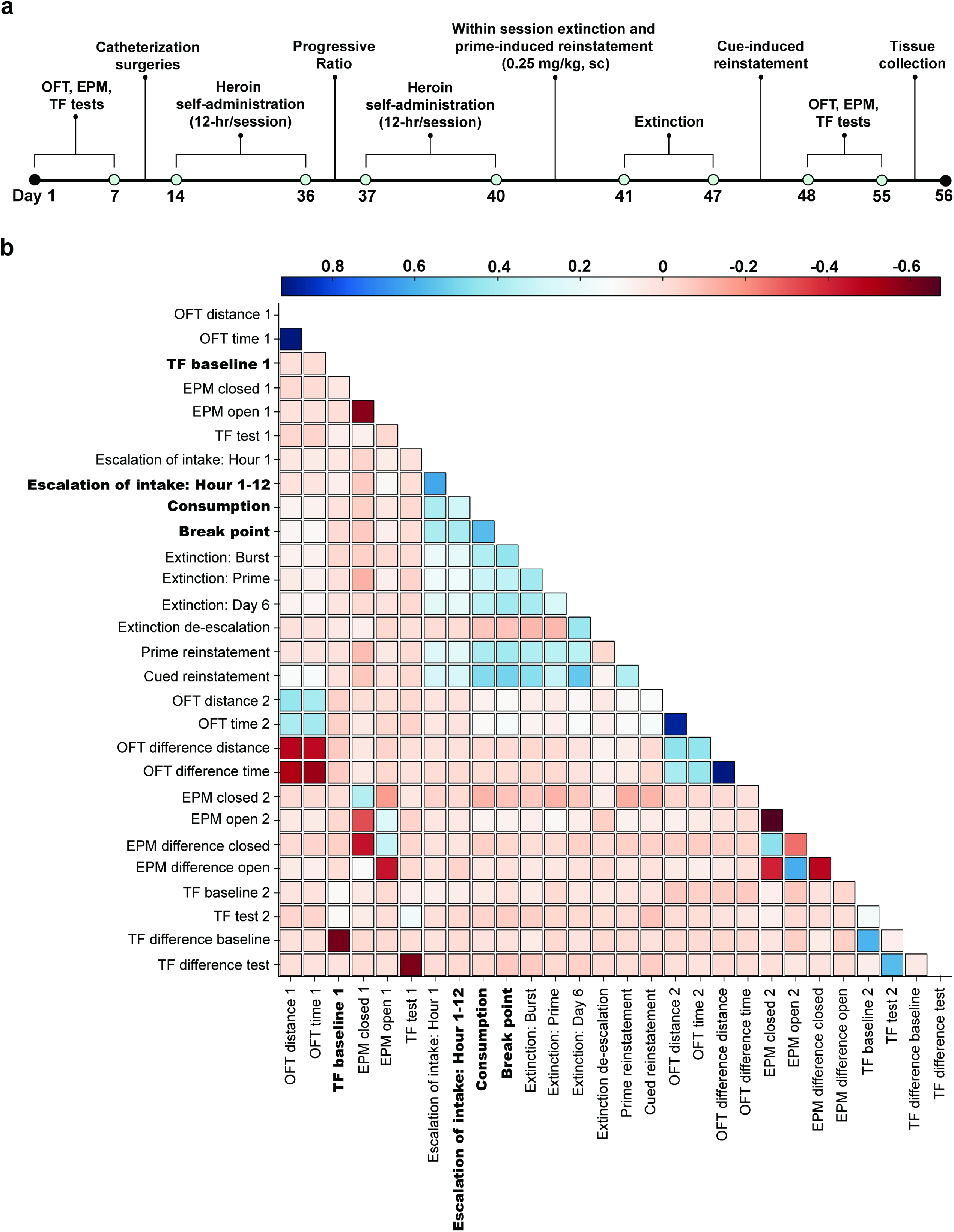
Experimental timeline and correlational analyses. **a)** Rats underwent testing for stress- and anxiety-related behaviors (OFT: open field test; EPM: elevated-plus maze) and nociceptive threshold (TF: tail flick) prior to jugular catheterization surgery, with tests repeated following heroin experience. Heroin self-administration occurred for 3 weeks (12-h/session, 4 days a week) followed by a progressive ratio test and an additional 3 days of self-administration training. Next, rats underwent a within-session extinction-heroin prime reinstatement test (6-h). Extinction training sessions (2-h/session) then occurred followed by a cue-induced reinstatement test (2-h). Spleen sample were collected for genetic analyses at the conclusion of training. **b)** Correlation matrix for behavioral trait covariance. Behavioral testing that occurred prior to heroin experience is designated with a 1, and testing following heroin training as 2. Traits in bold are those with at least one significant QTL identified in GWAS analysis. (n=874)

### Stress, anxiety and nociceptive-like behavioral measures

Stress and anxiety-related behaviors were assessed in the elevated-plus maze (EPM) and the open field test (OFT). Nociception under baseline (administration of 1 mg/kg saline, sc) and test (0.75 mg/kg heroin, sc) conditions were evaluated using the tail flick (TF) test. All tests were performed prior to and following heroin self-administration (see Supplemental Information and Kuhn et al., 2022 and 2024).

### Heroin taking, extinction and seeking measures

After EPM, OFT and TF tests, rats were outfitted with an indwelling jugular catheter prior to undergoing heroin self-administration training procedures. During training, presses on the active lever resulted in presentation of a tone and cue light stimulus, as well an infusion of heroin (20 µg/kg/100 µl infusion over 3-sec). Training occurred for 3 weeks with four 12-hr sessions/week. Next, motivation to take heroin was assessed using a progressive ratio test whereupon the number of lever presses needed to earn an infusion exponentially increased according to the following formula: (5*e^0.2n^)-5 (29). The maximum number of lever presses exerted to receive an infusion was considered the break point. Heroin self-administration training was then re-established for 3 days, followed by a within-session extinction-prime test lasting 6 hours. Rats were in extinction training conditions (i.e., no cue presentation or heroin delivery) for the entire 6-h session. However, with two hours left in the session, animals received a heroin prime injection (0.25 mg/kg, sc; heroin primed reinstatement test). Animals next underwent 6 daily 2-h extinction training sessions prior to a 2-h test for cue-induced reinstatement (i.e., active lever presses resulted in cue presentation but no heroin infusion). The EPM, OFT and TF tests were then repeated. See Supplemental Information for further details.

### Tissue collection and genotyping

Following all behavioral testing, animals were sacrificed and spleen samples were collected and shipped to University of California San Diego for genotyping. Genotyping-by-sequencing was employed as previously described to assess genotypes (30, 31). A total of 7,955,513 single nucleotide polymorphisms (SNPs) were identified with an error rate <1%. However, prior to GWAS analysis, the SNPs were filtered for missingness not above 10%, minor allele frequency (MAF) of no less than 0.5% and the Hardy-Weinberg equilibrium (HWE) deviation p-value no less than 1e^−10^, resulting in a final set of 5,308,726 SNPs for analysis.

### Statistical analyses

#### Behavioral traits assessed in GWAS

Behaviors included in GWAS analysis included behavioral testing traits (i.e., EPM, OFT and TF) both prior to (time point 1) and following (time point 2) heroin experience, as well as the difference in individual testing measures between the time points (Supplemental Table 1). Traits representative of heroin taking, extinction and seeking measures were also analyzed (Supplemental Table 1). Heroin-taking behaviors included: escalation of intake in the first hour of the session across training (ug/kg; average consumption hour 1 days 1-3 subtracted from hour 1 days 10-12; Escalation of intake: Hour 1), escalation of intake across the entire session (ug/kg; average consumption days 1-3 subtracted from days 10-12; Escalation of intake: Hour 1-12), total consumption (ug/kg) across training sessions 1-12, and break point achieved during the progressive ratio test. Extinction traits include active lever presses made during the following: hours 1-2 (Extinction: Burst) and 3-4 (Extinction: Prime) of the 6-h within-session extinction-heroin prime reinstatement test, the last day of extinction training prior to the test for cued reinstatement (Extinction: Day 6) and the difference in lever presses between the first and last day of extinction training (Extinction de-escalation). Active lever presses made during the tests for heroin prime and cued reinstatement were assessed for seeking measures.

#### Behavioral clustering

In addition to specific traits, we wanted to determine if any genetic variants were associated with OUD susceptibility. To do so, we employed a non-linear network-based clustering approach (32) using an R package ‘mlsbm’ (33) and characterized rats into OUD vulnerable, intermediate or resilient clusters. Behaviors included for analysis are indicated in Supplemental Table 1. Briefly, behavioral data was standardized within sex and site and combined to create a rat-rat similarity network, whereupon a Bayesian stochastic block model was employed to define clusters assignments (32). Behavioral and neurobiological validation of this model are extensively detailed in prior publications (23, 24). Cluster assignment was used as phenotypes (Vulnerable vs intermediate/resilient; Intermediate vs vulnerable/resilient; Resilient vs vulnerable/intermediate) for GWAS. Differences in cluster composition between the top SNP allelic variants were assessed using a Chi-square test.

#### Phenotypic correlations

To address difference in data variance between sites (MUSC vs UCAM), covariates (including sex) that explained more than 2% of trait variance (Supplemental Table 2 and 3) were regressed out and data was quantile normalized within site. This resulted in trait variances being equal between sites, thus weighing data from MUSC and UCAM equally. Next, data from both MUSC and UCAM were merged and quantile normalized again to maintain normality. Behavioral co-variance between traits was assessed using Spearman correlations with Bonferroni adjustment to correct for multiple comparisons using GraphPad Prism version 10. SNP heritability for behavioral traits was determined using GCTA-GREML analysis with significance determined by the p-value of rejecting the null hypothesis that the heritability is zero (p<0.05).

#### Quantitative trait loci (QTL) and SNP identification

To perform the GWAS we used GCTA software employing a genetic relatedness matrix and the Leave One Chromosome Out method to account for proximal contamination (34, 35). Because all data was quantile normalized, only one significance threshold, which was determined by permutation, was necessary (36). We report QTLs that surpassed the permutation-derived threshold of –log_10_ p>5.58 (p<0.05). Significant QTLs are only reported when there was at least one additional SNP located within 0.5 Mb with a significance within 2 –log_10_(p) units of the first SNP (37). Independence of QTLs on the same chromosome was established by making the top SNP in the most significant QTL a covariate and then repeating the GWAS analysis for the chromosome in question. This process was repeated until no more significant QTLs were identified (37). Linkage disequilibrium (LD; r^2^≥0.6) was used to define the boundaries of each QTL. SNPEff was used to identify coding SNPs within each QTL interval (38). To identify associations from other GWAS that have been performed in HS rats that map to the same loci, a phenome wide association study (PheWAS) was conducted. To explore functional mechanisms whereby SNPs may affect OUD-like traits, we assessed the co-location of the peak SNP in each QTL with known SNPs that can alter gene expression (eQTL), slicing patterns (sQTL) or directly alter the amino acid sequence of genes within a 3 Mb window using RatGTEx.org (28). The reference database for eQTLs was comprised of animals not in this study, therefore, simple correlations between expression and behavior were not possible. Importantly, the absence of coding variants, eQTLs or sQTLs could reflect the fact that RNA sequencing for the eQTL/sQTL analysis used a polyA pulldown protocol, limiting our findings to RNAs that are polyadenylated.

#### Principal component analysis

To assess whether variance in behavioral traits within significant QTLs could be explained by common factors, a principal component analysis was applied to the data. Number of components was determined using a minimum threshold of Eigenvalue of 1 and Varimax method with a minimum factor loading of 0.3 was used to rotate identified components. Analyses used SPSS version 27.

## Results

### Behavioral correlations

Behavioral traits showed weak to moderate correlations overall (Fig. 1b, Supplemental Table 4). However, behavior during the progressive ratio test, cued reinstatement and total heroin consumption were positively correlated (r^2^≥0.48), suggesting overlapping behavioral mechanisms guiding these behaviors.

### Heritability estimates

SNP heritability (h^2^) estimates the proportion of the phenotypic variance explained by the genetic variance and is dependent on the genetic diversity of the population and environmental sources of variability. Aspects of heroin taking (Consumption, h^2^=0.17) and the inability to initially refrain from seeking (Extinction: Burst, h^2^=0.10) were heritable (Fig. 2a, Supplemental Table 5). While some measures of heroin taking, extinction and seeking were not heritable, OUD susceptibility cluster assignment (23, 24) was heritable (Vulnerable vs resilient/intermediate, h^2^=0.17; Intermediate vs vulnerable/resilient, h^2^=0.09; Resilient vs vulnerable/intermediate, h^2^=0.15), suggesting a heritable genetic contribution to OUD susceptibility (Fig. 2a, Supplemental Table 5).

**Figure 2.**
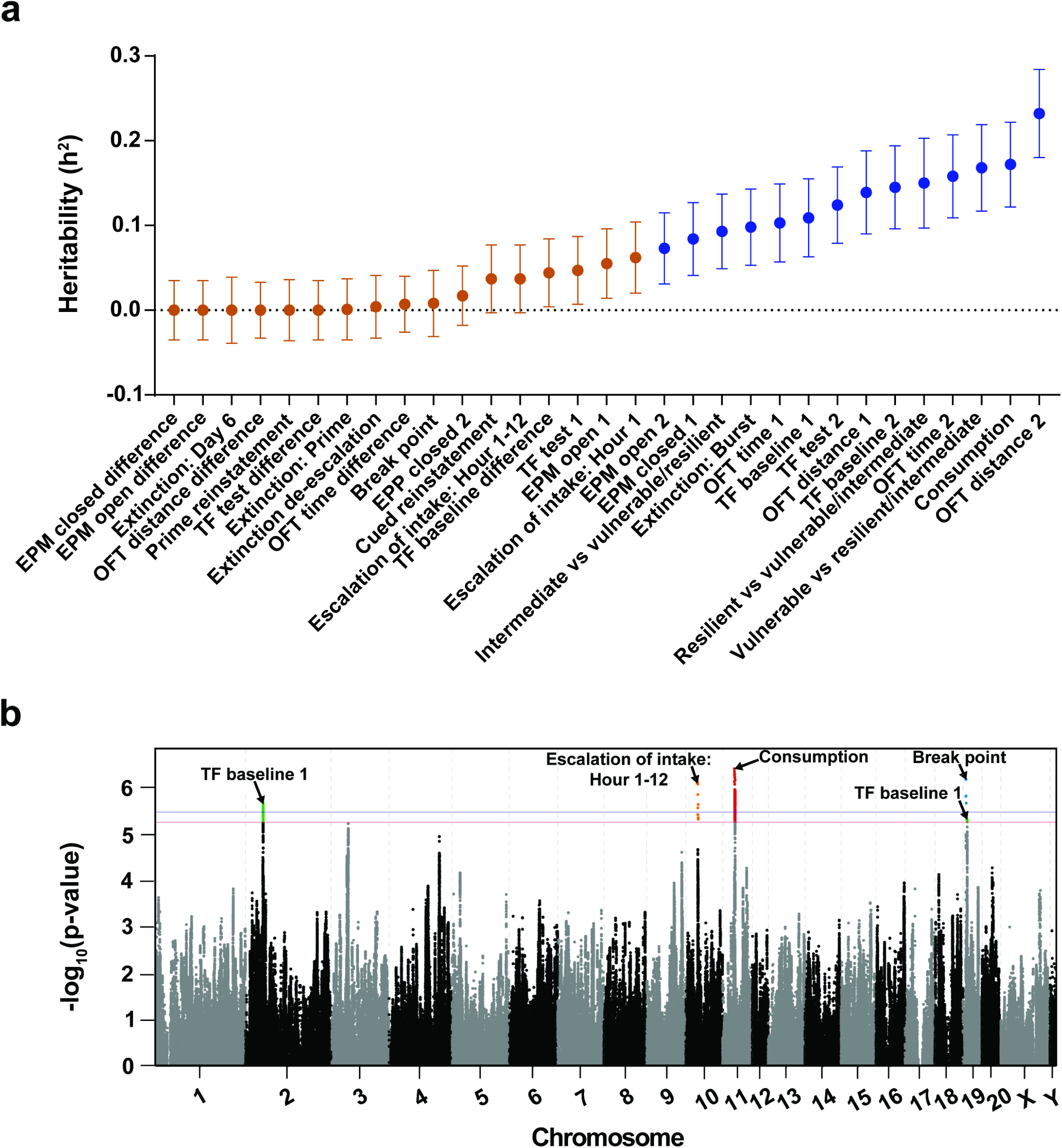
Heritability and genome-wide association results. **a)** SNP heritability (h^2^) measure ± SEM for all behavioral traits assessed. Significant h^2^ measures (p<0.05) are indicated in blue, and non-significant traits are in orange. **b)** Porcupine plot for selected behavioral traits. Top SNPs exceeding the genome-wide significance threshold are highlighted and identified (red: −log(p)=5.36 and blue: −log(p)=5.58, which correspond to permutation derived p values of 0.10 and 0.05, respectively. (n=874)

Several behavioral measures prior to and following heroin administration experience showed significant heritability (Fig. 2a, Supplemental Table 5). Nociceptive threshold under basal conditions both prior to (TF baseline 1, h^2^=0.11) and following (TF baseline 2, h^2^=0.15) heroin experience were both heritable. Heroin-induced nociception after prolonged heroin exposure was also heritable (TF test 2, h^2^=0.12), reflecting the genetic contribution to the long-term effects of heroin exposure on nociceptive threshold. Aspects of anxiety- and stress-like behaviors were also heritable, including high anxiety prior to heroin experience (EPM closed 1, h^2^=0.08), but lower levels following heroin self-administration (EPM open 2, h^2^=0.07). Our results support previous findings that locomotor behavior is heritable (25, 26) (OFT distance 1, h^2^=0.14; OFT time 1, h^2^=0.10) and show that this heritability persists even after exposure to heroin administration (OFT distance 2, h^2^=0.23; OFT time 2, h^2^=0.16).

### GWAS

We identified genome-wide significant associations for several traits: escalation of intake across the entire session on Chromosome 10, total heroin consumption on Chromosome 11, break point achieved during the progressive ratio test on Chromosome 19, and baseline nociception prior to heroin experience (TF baseline 1) on Chromosomes 2 and 19 (Fig. 2b). Shared behavioral variance between these traits was assessed using PCA, with heroin taking traits loading onto one factor and baseline nociception onto its own factor, reflecting the independence of these measures in our data (Supplemental Fig. 1). While both nociception and break point were associated with loci on Chromosome 19, the associated loci are separated by approximately 4 Mb and are not in strong LD with one another, indicating that they are due to separate causal loci. See Supplemental Table 6 for MAF and HWE values for each significant loci and the Supplementary Information for a detailed report of all GWAS findings. Both sites contributed comparably to significant loci (Supplemental Table 7).

*Escalation of intake:* We identified a significant locus on Chromosome 10 for escalation of heroin intake (Fig. 3). The peak SNP was located at 10:35,034,818 and spans from positions 34,961,844 to 35,103,415. This locus did not contain any coding variants that are in strong LD with the top SNP. Though eQTLs and sQTLs were identified, none are in strong LD (i.e., r^2^) with the top SNP, suggesting they are unlikely to be causally related to the escalation of heroin intake behavior. Identified eQTLs include metabotropic glutamate receptor 6 (*Grm6*) in whole brain (r^2^=0.63), collagen type XXIII alpha 1 chain (*Col23a1*) in the orbitofrontal cortex (OFC; r^2^=0.63) and zinc finger protein (*Zfp2*) in the prelimbic cortex (PrL; r^2^=0.64). Two sQTLs were also identified, jade family PHD finger 2 (*Jade2*) in the nucleus accumbens core (NAcc) and basolateral amygdala (BLA; both r^2^=0.63), and *Col23a1* in the PrL (r^2^=0.63). The lack of any putatively causal variants in this interval suggests that the causal variant remains unknown, possibly because there is an eQTL or sQTL in a tissue that is not represented in the RatGTEx database, or an eQTL that was not detected, for example, because it is significant at a different developmental time point or under different environmental conditions.

**Figure 3.**
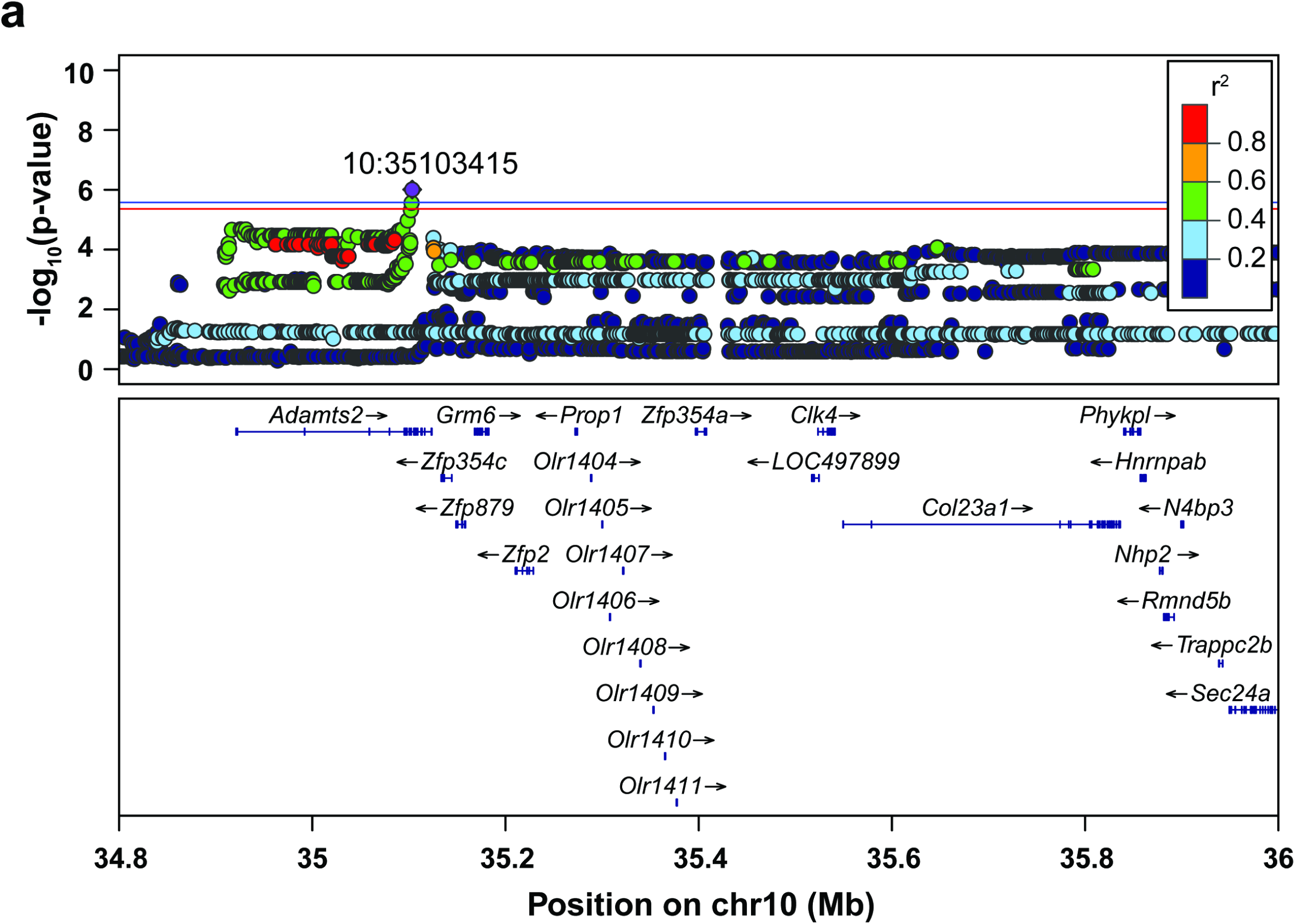
GWAS results for escalation of heroin intake across the entire session. Regional association plot for chromosome 10, with the top SNP in purple and chromosome location indicated on the plot. Other SNPs are colored based on their linkage disequilibrium with the top SNP. Known genes of SNPs are indicated below the plots. Significance threshold: red line, p<0.10; blue line, p<0.05. (n=874)

*Total consumption:* Total heroin consumed across training (Supplemental Fig. 2) was associated with a locus on Chromosome 11 (Fig. 4a) that spanned from positions 35,033,985 to 36,454,941 with the peak SNP at 35,034,818. Several significant causal coding variants were identified that are in strong LD with this locus, including bromodomain and WD repeat domain containing 1 (*Brwd1*: Leu2187Pro, r^2^=0.99), Purkinje cell protein 4 (*Pcp4*: Lys12Arg, r^2^=0.99) and two different amino acid coding variants for SH3 domain binding glutamate-rich protein (*Sh3bgr:* Glu169Lys, r^2^=0.99; Glu177Lys, r^2^=0.99). There were also eQTLs for *Sh3bgr* that was in high LD with this locus (r^2^=0.99 for all) in whole brain, BLA, NAcc, PrL, lateral habenula (LHb) and OFC. Two additional genes with eQTLs in high LD with the locus were identified: *Pcp4* (PrL, r^2^=0.88) and a long non-coding RNA (*ENSRNOG00000068065*; infralimbic cortex (IL), r^2^=0.99). We also identified an sQTL for *ENSRNOG00000068065* in several brain regions (whole brain, BLA, LHb, OFC, NAcc, PrL, and PrL2; r^2^=0.99-1.00). Lastly, there was an sQTL for the gene guided entry of tail-anchored proteins factor 1 (*Get1*; r^2^=0.99) in the IL. Heroin consumption is likely mediated, in part, by these coding differences and or by heritable differences in expression of one or more of these genes. Because several plausible variants are in LD with one another, mechanistic or other complementary approaches will be needed to identify the causal gene. Additionally, PheWAS analysis showed that OUD vulnerability (Vulnerable vs Intermediate/resilient cluster) was associated with the top SNP at a suggestive level (-log(p)=4.5), with cluster composition differing between the allelic variants (x^2^=18.12, p<0.0001; Fig. 4b,c).

**Figure 4.**
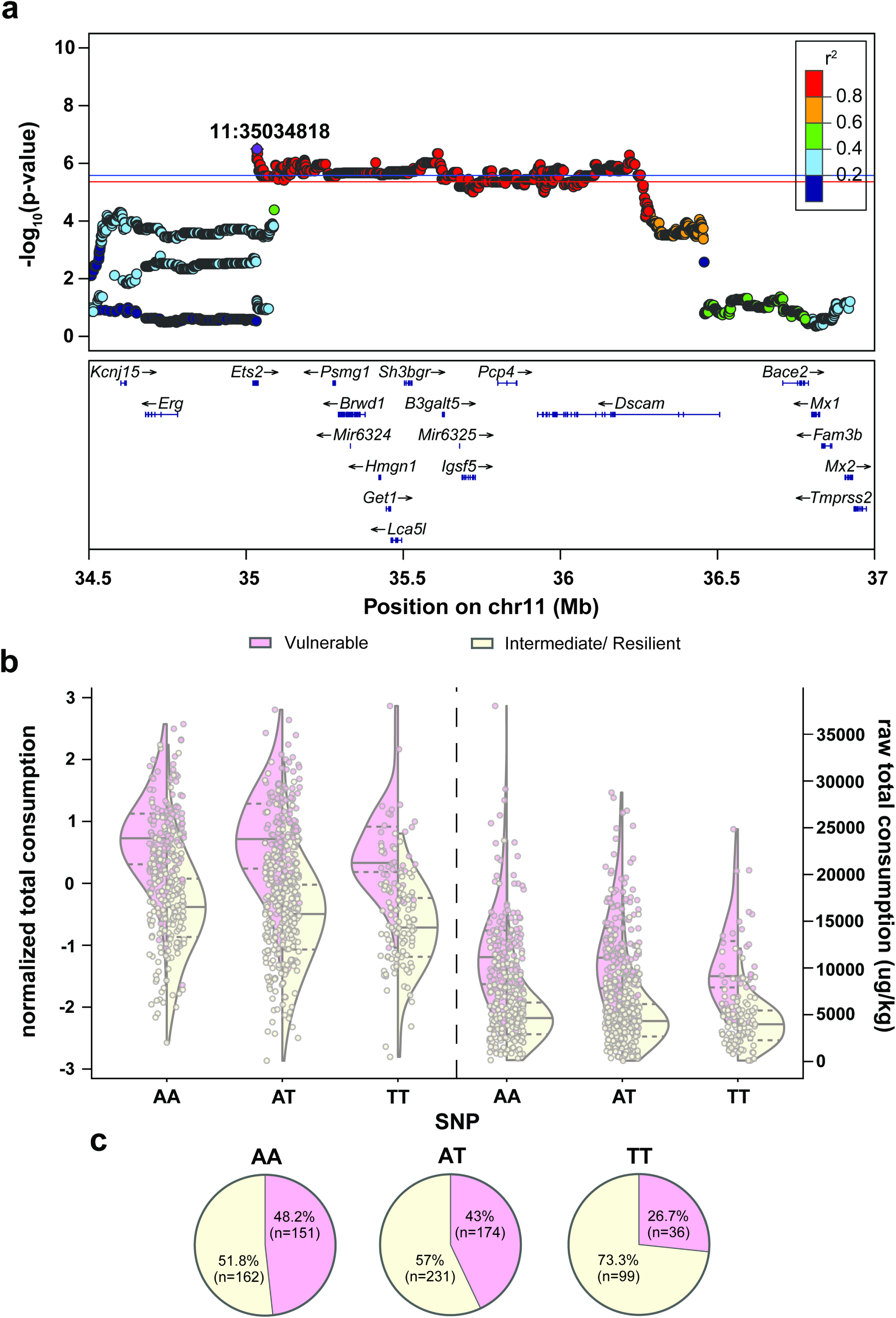
GWAS results for total heroin consumed across self-administration training. **a)** Regional association plot for chromosome 11, with the purple dot indicating the top SNP and location on chromosome indicated on the plot. The color of additional SNPS refers to extent of linkage disequilibrium with the top SNP. Known genes of SNPs are indicated below the plots. **b)** Effect plots for total heroin consumption during training (left: normalized data; right: raw data) for OUD phenotypes (Vulnerable: pink; Intermediate/resilient: yellow) according to peak SNP allelic variant. Half of a violin plot for each phenotype is shown, with the solid line indicating the median value of the data and the dash lines representing the upper and lower interquartile range. **c)** Top SNP allelic variant OUD phenotype cluster composition differ (x^2^(1, 853)=18.12, p<0.0001), with the minor allele associated with greater OUD resiliency. Significance threshold: red line, p<0.10; blue line, p<0.05. (n=853; Vulnerable (pink), n= 361; Intermediate/resilient (yellow), n= 492)

*Break point:* The break point achieved during the progressive ratio test was associated with a locus on Chromosome 19 (Fig, 5a) spanning from positions 8,647,094 to 8,801,084 with the peak SNP at 8,648,697. Two potentially causal coding variants in prohibitin 1 like 2 (*Phb1l2*; Lys70Asn and Ile231Asn), which were in perfect LD with the top SNP (r^2^=1.0), were identified. An eQTL with perfect LD with the locus for the gene matrix metallopeptidase 15 (*Mmp15*; r^2^=1.0) was also identified in the whole brain. An additional eQTL (solute carrier family 38 member 7, *Slc38a7*) and sQTL (casein kinase 1 alpha 2, *Csnk2a2*) were identified in the BLA, however, the relationship between these genes and the locus are modest (r^2^=0.65 for both) rendering them much less likely to be the causes of the association with break point behavior. Similar to the Chromosome 11 association for total heroin consumption, PheWAS analysis showed a suggestive association between the top SNP and the OUD vulnerable cluster (Vulnerable vs intermediate/resilient clusters; −log(p)=5.16; Fig. 5b). OUD cluster composition differed between the alleles at the peak SNP (x^2^=18.91, p<0.0001; Fig. 5c). While there is a small bias toward the AA variant conferring OUD resiliency, the minor variant, AC, strongly coincides with OUD vulnerability.

**Figure 5.**
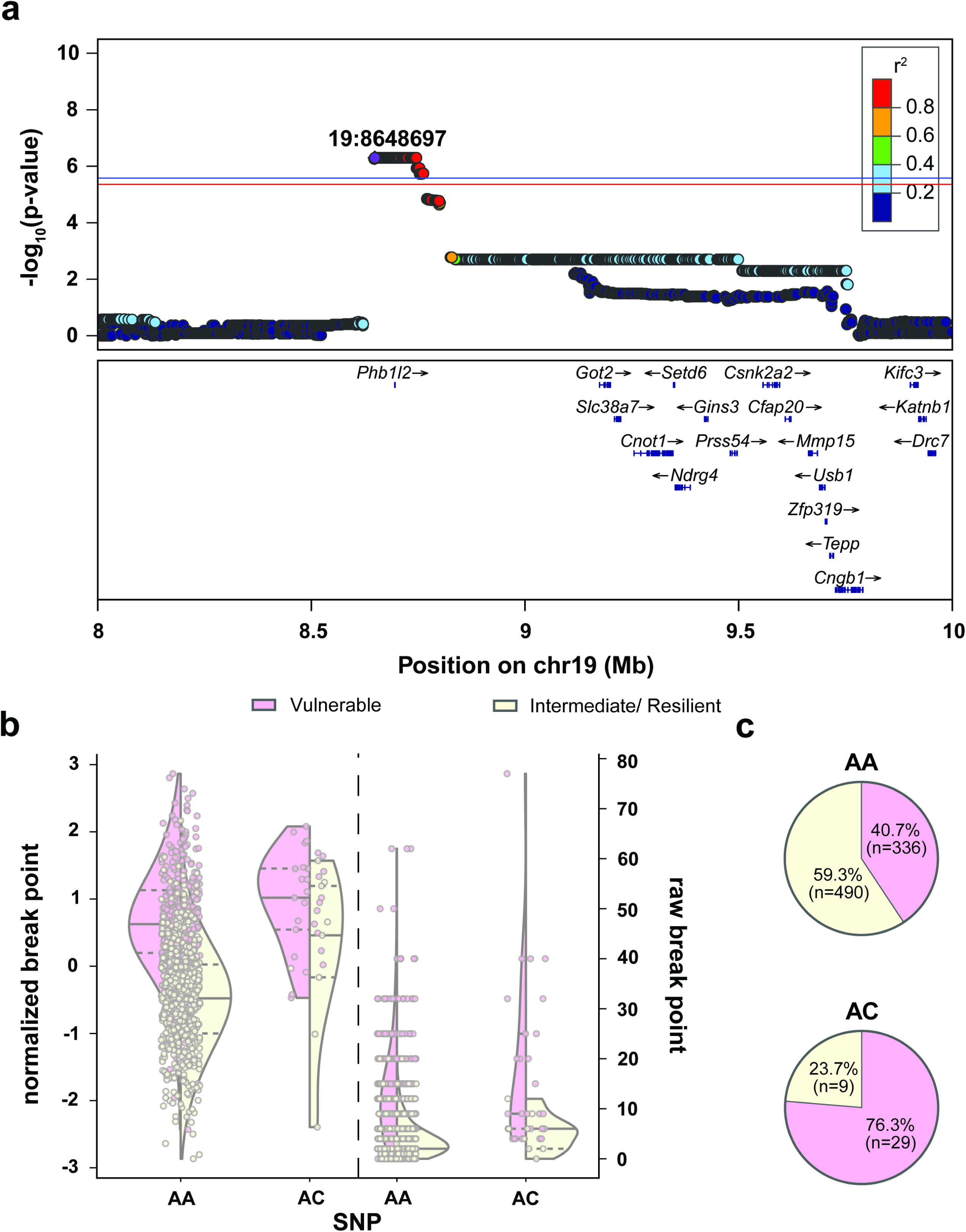
GWAS results for break point achieved during the progressive ratio test. **a)** Regional association plot for chromosome 19, with the top SNP in purple and the color of other significant SNPs referring to level of linkage disequilibrium with the top SNPs. The location of the top SNP is indicated on the plot. Known genes of SNPs are indicated below the plots. **b)** Effect plots showing break point achieved during the progressive ratio test (left: normalized data, right: raw data) for OUD vulnerable (pink) and intermediate/resilient (yellow) rats in relation to the top SNP allelic variant. Half a violin plot is shown for each OUD phenotype, with the solid line indicating the median value of the data set, whereas the dashed lines represent the upper and lower interquartile range. **c)** OUD phenotypic cluster composition for both peak SNP allelic variants. Cluster composition between variants differed (x^2^(1, 864)=18.91, p<0.0001), with the minor allele heavily biased toward an OUD vulnerable phenotype. Significance threshold: red line, p<0.10; blue line, p<0.05. (n=864; Vulnerable (pink), n= 345; Intermediate/resilient (yellow), n= 519)

*Baseline nociception prior to heroin experience:* A locus on Chromosome 2 spanning positions 51,450,191 to 52,982,078 with the peak SNP at 51,588,876 was identified for basal nociception prior to heroin taking procedures (Fig 6a). Several coding variants in modest LD with the peak SNP were identified in genes within this locus, including coiled-coil domain containing 152 (*Ccdc152*: Ala242Ser and Thr14Asn, both r^2^=0.77), growth hormone receptor (*Ghr*: Ala546Val and Lys541Asn, both r^2^=0.77), zinc finger protein 131 (*Zfp131*: Gly506Ser, r^2^=0.82) and serine/threonine-protein kinase NIM1 (*Nim1k;* r^2^=0.64). The gene selenoprotein P (*Selenop*) had an sQTL in both BLA and whole brain (r^2^=0.78-0.79), and an eQTL for both whole brain and the NAcc (both r^2^=0.66). These data provide support of the involvement of these genes in the Chromosome 2 association with baseline nociception.

**Figure 6.**
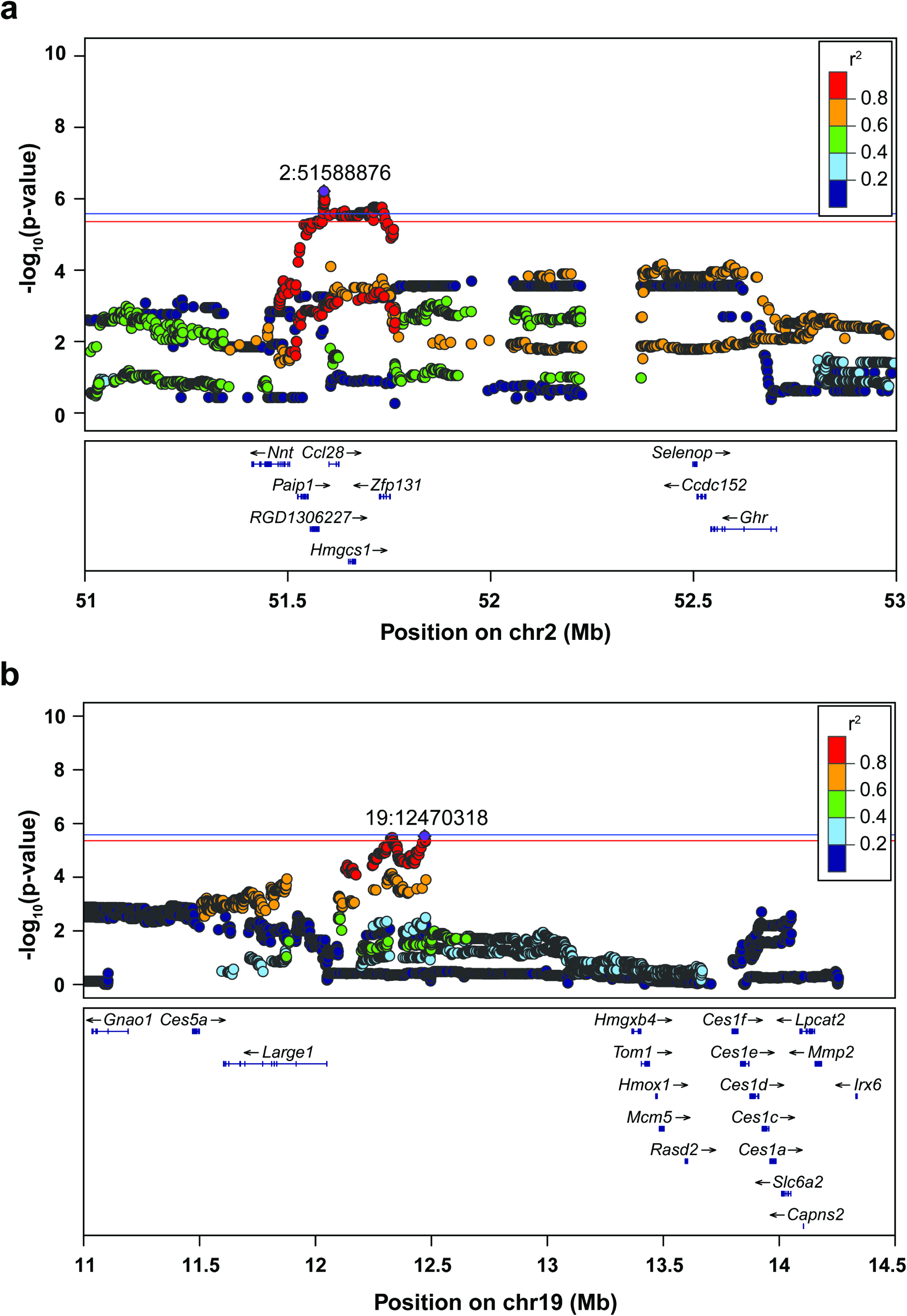
GWAS results for analgesic threshold during the TF baseline session prior to heroin experience. Regional association plots for **a)** chromosome 2 and **b)** chromosome 19. The top SNP identified for each chromosome is in purple with chromosome location indicated on the plot. The color of other SNPs corresponds to extent of linkage disequilibrium with the top SNP. Known genes of SNPs are indicated below the plots. Significance threshold: red line, p<0.10; blue line, p<0.05. (n= 874)

An additional significant association for this behavior was identified on Chromosome 19 (Fig. 6b). The peak SNP was located at 19:12,470,318 with the QTL spanning from positions 11,508,674 to 12,474,455. A potentially causal coding variant in modest LD with the top SNP in the gene LARGE xylosyl- and glucuronyltransferase 1 *(Large1*: Ala92Val, r^2^=0.75) was present. There were also several eQTLs for *Large1* (whole brain, r^2^=0.69; BLA, r^2^=0.78; IL, r^2^=0.68; NAcc, r^2^=0.70; OFC, r^2^=0.69; PrL, r^2^=0.70; and PrL2, r^2^=0.75). In addition, there was an eQTL for ribosomal protein S18 (*Rps18I2*) within the LHb (r^2^=0.98). An sQTL for the gene myb1 trafficking protein (*Tom1*) was in strong LD with this locus (r^2^=1.0) within the PrL. Finally, there was a significant sQTL for metallothionein 3 (*Mt3*) in the BLA (r^2^=0.75). These coding variants, eQTLs and sQTLs could point to heritable differences and underlying biological mechanisms for basal nociception prior to drug experience at the Chromosome 19 locus.

## Discussion

The economic toll on society for OUD totals over one trillion dollars annually (39). Furthermore, over one million individuals seek treatment for OUD annually (40) and more than 80,000 people die from opioid overdoses every year in the US alone (41). Thus, there is an urgent need to understand the factors that influence vulnerability to OUD. Using a rat line emulating within species genetic and phenotypic variability (14) akin to humans, we assessed the genetic basis of several OUD-like behaviors and further extended analysis to account for OUD susceptibility. While heroin was used, we expect these findings will be relevant for other opioids of abuse whose primary site of action is mu opioid receptors. We found several traits related to OUD are heritable, including overall OUD susceptibility. Genetic variants associated with nociception prior to heroin experience as well as heroin consummatory behaviors were identified, several of which are either biomarkers for other SUDs or are involved in neuroplasticity. Using this model, we are able to identify genetic variants possibly associated with an OUD diagnosis, but also specific behavioral traits that comprise vulnerability or resiliency thereby further contributing to our knowledge of the genetic basis of OUD.

### Heritability

OUD is heritable in humans and we found several OUD-like behaviors were also heritable in HS rats. Human twin studies have yielded heritability estimates between 23-54% for OUD (4, 5) and human GWAS of OUD have produced SNP heritability estimates, which are expected to be lower, ranging from 6-12.7% (9–13). However, given the complex and multi-symptomatic nature of OUD (42), assessing heritability of distinct behavioral traits associated with human OUD vulnerability is difficult. In a large sample of HS rats, we identified several heritable heroin addiction-like traits that model different aspects of human OUD. Both heroin consumption and initial extinction behavior were significantly heritable, with SNP heritability estimates for these traits similar to SNP heritability obtained from human studies of OUD. However, behaviors following prolonged heroin use and forced abstinence, such as cued reinstatement, did not exhibit significant heritability, suggesting weaker genetics influences and/or limitations in our procedure’s ability to detect the cascade of neuroplastic adaptations that occur following prolonged heroin self-administration (43, 44).

Novelty-induced locomotion, a heritable behavior (25, 26), has been associated with vulnerability in opioid addiction-related behaviors in both rodent models (17, 45–49) and humans (50). We confirmed the heritability of locomotor behavior both prior to and following heroin experience in HS rats, suggesting an enduring genetic contribution to this behavior that remains relatively unaffected by factors (e.g., epigenetic changes) associated with prolonged heroin experience. Lastly, in agreement with prior work in humans, we showed that nociceptive threshold to heat stimuli under baseline conditions is heritable (51). Following opioid use, both humans (52, 53) and rats (54) demonstrate opioid-induced hyperalgesia in response to a painful stimuli (17), a finding replicated in our study. However, the heritability of nociception in humans is unknown. Given the inherent behavioral and genetic heterogeneity present in HS rats, and the translational relevance of the findings reported here, HS rats offer an opportunity to disentangle the genetic factors contributing to opioid-induced nociception.

### GWAS

Genetic variants associated with OUD-like behaviors in rats can contribute novel insights that improve our understanding of the neurobiological mechanisms underlying distinct behavioral traits associated with OUD in humans. This study implicated several loci and genes within those loci that had coding variants, eQTLs and/or sQTLs associated with affective or behavioral traits associated with SUD (Supplemental Table 8).

#### Escalation of heroin intake across the entire session

While no causal coding variants, eQTLs or sQTLs were in strong LD with the QTL on Chromosome 10 for escalation of heroin-taking behavior across the entire session, several genes in this interval are of potential interest. For example, *Adamts2*, an extracellular matrix proteinase (55) localized within the mesolimbic and mesocortical regions within the brain (56), is a biomarker for heavy tobacco smoking and is downregulated following long-term morphine exposure and withdrawal (57). Similarly, the gene heterogeneous nuclear ribonucleoprotein H1 (*Hnrnph1*) is located in this interval. *Hnrnph1* was identified as the causal gene for a mouse QTL for methamphetamine response (58–61) and subsequent work has shown it also modulates the response to fentanyl (62) and alcohol (63). Further investigation is required to determine if there are overlooked coding variants or eQTLs for *Adamts2* or *Hnrnph1*.

#### Heroin consumption

The region on Chromosome 11 that was associated with total consumption of heroin across all training sessions contained a number of genes with coding and expression variants. Three genes harbored coding variants that were in strong LD with the peak SNP: *Brwd1*, *Pcp4* and *Sh3bgr*. In humans, *BRWD1* has been associated with potentially relevant traits including cigarette smoking (64–69), externalizing behaviors (70), ADHD (67, 71) and hyperthymia in bipolar disorder (72). *PCP4* mediates calcium signaling and through downstream mechanisms also affects dopamine and acetylcholine signaling [76]. *PCP4* levels are attenuated in post-mortem tissue of individuals diagnosed with alcohol use disorder [77] as well as in the NAc of rats following 2-weeks of nicotine withdrawal [78]. This gene has also been linked to cigarette smoking (64, 73) and peripheral effects of alcohol drinking (74) in humans. We identified an eQTL for *Pcp4* in the PrL, a critical component of drug-reward circuitry and plasticity (75, 76). The relationship between *PCP4* and OUD is unknown, but our results suggest it may act to alter cortical to NAc transmission as a consequence of sustained heroin use. Finally, *Sh3bgr* had two coding variants and numerous eQTLs in whole brain, BLA, NAcc, OFC, LHb, and PrL, however the homologous human gene has not been associated with any psychiatric traits in prior human GWAS. In addition to these genes, a long non-coding RNA (ENSRNOG00000068065) and the gene *Get1* had sQTLs, but there is no supporting literature for their role in OUD-relevant behaviors.

*Ets2*, which contained the peak SNP for this locus, did not identify any coding variants or an e/sQTL. *Ets2* encodes a ubiquitous transcription factor that is found in many cells and has be linked to molecular processes (77) and cell types (78) that modulate synaptic remodeling, including those associated with OUD (79). Our results suggest *Ets2* and its allelic variants may impact OUD susceptibility, though further investigation is necessary.

While the functional association between heroin consumption and the other identified e/sQTLs are not known, *DSCAM* is correlated with the top SNP and encodes a neuronal cell adhesion molecule that regulates cellular connectivity (80) and glutamatergic plasticity associated with learning (81, 82). Glutamate-mediated neuroplasticity occurs following heroin experience and is considered a driving force behind relapse vulnerability (44). *DSCAM* is necessary for tasks requiring memory formation (83, 84), including those associated with rewarded behaviors (84), and was identified in human GWAS as a biomarker for cigarette cessation success (85). Based on these data, *DSCAM* warrants further investigation for OUD.

#### Break point

The QTL on Chromosome 19 for break point achieved during the progressive ratio test contained two coding variants in strong LD with the top variant for the poorly characterized gene *Phb1l2*. *Phb1I2* is localized to both neuronal and non-neural cell types in the central nervous system and helps to maintain mitochondria respiration by attenuating reactive oxygen species which contribute to cellular dysfunction (86). Chronic morphine administration disrupts mitochondrial respiration and restoring proper functioning attenuates opioid withdrawal (87). Additionally, *Phb1*, an analogue of *Phb1l2*, has been implicated in mediating glutamatergic and dopaminergic neural transmission and becomes dysregulated in several disorders, including alcohol use disorder (86). It is thus possible that differences in the coding sequence of *Phb1l2* influence break point through glutamate and dopamine dependent processes. Additionally, an eQTL for *Mmp15* in whole brain that was in strong LD with the top SNP was identified, suggesting *Mmp15* expression could modulate break point. MMP15 activates MMP2 (88) which contributes to NAcc cell-specific regulation of extinction behavior following heroin self-administration (89). It is possible that attenuated *Mmp15*-induced *Mmp2* expression contributes to higher break point levels. Furthermore, PheWAS analysis identified this locus as being associated with OUD vulnerability with the coding variant likely impacting OUD susceptibility.

Several of the heroin taking traits are both phenotypically and genetically correlated. Despite this, the significant loci identified for these traits are not the same (Supplemental Table 9). We attribute this observation to the fact that these traits are highly polygenic, meaning that many loci make small contributions to the observed phenotype. Because our power is limited, we only expect to identify a small subset of the true positive loci, and those subsets are expected to differ from one trait to another for purely stochastic reasons.

#### Baseline nociceptive threshold prior to heroin experience

Genetic variants on two separate chromosomes were found for nociceptive threshold under baseline conditions, prior to heroin experience. On Chromosome 19, *Large1* was notable because it contained both putatively causal coding variants and an eQTL. Large1 has been associated with various behavioral traits in humans, including educational attainment (90–92), smoking (64), alcohol dependence (93), neuroticism (94) and depression (94, 95); however, no prior evidence that we are aware of link it to nociception. Heritable differences in splicing patterns (sQTLs) for both *Tom1* in the PrL and *Mt3* in the BLA are also candidates for nociception, though neither has been previously directly linked to pain processing. *Mt3* is a brain zinc binding protein involved in several metabolic processes and has been implicated in cellular dysfunction of diseases including Alzheimer and Huntington disease (96). *Tom1* is an integral component of microglial function (97), which is involved in neuropathic pain (98), but has yet to be implicated in acute painful stimuli. This gene has also been associated with cigarette smoking in humans (73).

On chromosome 2, two genes that operate in tandem with one another were also associated with nociception. There was modest LD between the top SNP for this QTL and splicing patterns in *Selenop* which encodes for a selenoprotein. This protein is secreted by the liver and acts to deliver selenium, a trace element necessary for proper brain function (99), via the bloodstream to regions including the brain. Selenium has been implicated in mediating dopamine release and transmission within the substantia nigra (100, 101) and the NAc (102). The presence of *Selenop* on NAc dopamine terminals is proposed to attenuate dopamine release into the synaptic cleft via direct mediation of dopamine 2 autoreceptors, protecting against neurotoxic effects of drugs of abuse such as methamphetamine (102). *Ccdc152*, which also had coding variants that were in modest LD with this QTL, is encoded within the *Selenop* gene, and acts to attenuate *Selenop* levels via post-translational modification (103). Though the periaqueductal gray is the predominant brain nuclei processing pain signaling (104), striatal dopamine signaling has recently been implicated in affecting pain processing and perception (105, 106) and warrants further investigation.

## Conclusion and Future Directions

We used a rodent model to identify several heritable traits and genetic variants associated with OUD. Unlike GWAS performed in humans, we were able to identify genetic variants associated with distinct OUD behaviors that contribute toward diagnosis. Several of the identified genes are supported by prior relationships to SUDs and other relevant behavioral and psychiatric traits, suggesting that they warrant further investigation for OUD. Furthermore, using an established rodent model that characterizes rats using a non-linear compilation of behaviors across the SUD phases to confer OUD susceptibility, we identified genetic variants for heroin consumption and motivation that were associated specifically with OUD vulnerability. These findings strengthen the translational utility of the behavioral model and can help to assess genetic variants and biomarkers associated with OUD vulnerability versus resiliency. However, complementing human OUD (5), phenotypic variance can only be partially explained by the genetic variants discovered here. Other genetic variants as well as epigenetic (107) and uncontrolled environmental factors also contribute to OUD behaviors in HS rats. Furthermore, both preclinical and clinical GWAS analysis focusing on genetic variants associated with OUD resiliency, a neurobiological active process (108, 109), should be considered in future analyses.

## Acknowledgements

The authors would like to thank the numerous technicians within the Kalivas and Ciccocioppo lab for their help performing the behavioral assays on the rats used in these analyses, as well as members of the Kalivas lab for their helpful feedback on the manuscript. This work was supported by the National Institute on Drug Abuse U01DA45300 (PWK), P50DA037844 (AAP), T32DA007288 (BNK) and K99DA057390 (BNK).

## Supplementary Information

### Behavioral testing

#### Elevated-plus maze (EPM)

Sessions were conducted in an elevated apparatus (San Diego Instruments; 4 arms at 110.5 cm long and 10.2 cm wide) that consisted of two “closed” arms (30.5 cm high walls) and two “open” arms (i.e., not enclosed). Animals were placed in the center of the maze to start, and movement was tracked automatically using ANY-maze behavioral tracking software (Stoelting, Wood Dale; version 6.17) over the course of the 5-min session. A minimum of 85% of the rat’s body had to be within an arm for it to be counted. The percent time the animal spent in each arm was calculated.

#### Open field test (OFT)

Animals were placed in the center of a Plexiglas chamber housed within a metal frame (Omnitech Electronics, Columbus, OH; 40.6 cm L x 40.6 cm W x 30.5 cm H) containing photocells that tracked vertical and horizontal movements. Sessions lasted for 60-min and data was recorded and analyzed for distance and time spent moving using Versamax (Omnitech Electronics, Columbus, OH; version 1.80-0142).

#### Tail flick test (TF)

Testing occurred on a platform with the rat’s tail over an infrared light beam equipped with a motion sensor that turned the beam off once the tail was removed, or after 10 seconds (Ugo Basile S.R.L., Gemonio, Italy). TF was comprised of two phases: baseline (1 mg/kg saline injection, s.c.) and test (0.75 mg/kg heroin, s.c.). Injections were administered 15 min prior to testing and the phases were separated by 1 hour. Each test consisted of 4 trials with the location of the tail tested being adjusted by 1 cm each trial to prevent tissue damage. Average latency to remove the tail from the beam was calculated.

### Heroin taking, refraining and seeking measures

Prior to training, rats were outfitted with an indwelling jugular catheter and post-operatively administered an antibiotic (Cefazolin, 0.2 mg/kg, s.c.; or enrofloxacin, 1 mg/kg, i.v.) and analgesic (Ketorolac, 2 mg/kg, s.c.; or Meloxicam, 0.5 mg/rat, s.c.). Training occurred in standard behavioral testing chambers (Med Associates, St. Albans, VT) housed within a sound attenuating box outfitted with a ventilation fan. Two levers with a light above each were on one chamber wall, and opposite was a house light and speaker. The house light turned on at the start of all sessions. A fixed ratio 1 schedule of reinforcement was used for heroin self-administration whereupon a press on the active lever resulted in an infusion of heroin (20 µg/kg/100 µl infusion over 3-sec) and a 5-sec presentation of a light/tone cue. The house light turned off for 20-sec at the time of infusion to signal a time out period during which additional active lever presses were recorded but without consequence. Four training sessions (12-h or 300 infusions earned) occurred per week (Monday-Friday) with one randomized day off per week, totaling 12 sessions. The progressive ratio test occurred next to assess heroin break point. Testing terminated after 12-h or 1-h of no earned infusion. Heroin self-administration training was then re-established for 3 days, followed by a within-session extinction-prime test lasting 6 hours. Testing occurred under extinction conditions (i.e., active lever presses no longer resulted in heroin infusion or light/tone cue presentation). With two hours left in the session, animals received a heroin prime injection (0.25 mg/kg, s.c.). Extinction training session and a test for cued reinstatement followed. Throughout testing, presses on the inactive lever were recorded but had no consequence.

### GWAS analysis

A detailed report of all GWAS findings may be found in the included ZIP file.

**Supplemental Table 1.** Description of behavioral traits assessed in GWAS. Traits included those measured both prior to (time point 1) and following (time point 2) heroin experience, and the difference between the two time points, along with several behaviors associated with heroin taking, refraining and seeking. Traits used in the non-linear clustering approach to categorize rats into OUD phenotypes are designated with an asterisk.

| Trait | Description |
| --- | --- |
| *Prior to heroin experience* |  |
| OFT distance 1 | Total distance travelled |
| OFT time 1 | Time spent moving |
| EPM closed 1 | Time spent in closed arms |
| EPM open 1 | Time spent in open arms |
| TF baseline 1 | Average latency of tail removal during baseline (i.e., saline) session |
| TF test 1 | Average latency of tail removal during test (i.e., heroin) session |
| *Heroin measures* |  |
| Escalation of intake: Hour 1 | Average consumption ($\mu\text{g/kg}$) from hour 1 for sessions 1-3 subtracted from average hour 1 consumption for sessions 10-12 |
| *Escalation of intake: Hour 1-12 | Average consumption ($\mu\text{g/kg}$) from entire session for sessions 1-3 subtracted from average total consumption for sessions 10-12 |
| *Consumption | Cumulative heroin consumption ($\mu\text{g/kg}$) for sessions 1-12 |
| *Breakpoint | Break point (i.e., most active lever presses willing to expend to receive an infusion of heroin) during progressive ratio test |
| *Extinction: Burst | Active lever presses during the first 2 hr of the within-session extinction-prime reinstatement test |
| Extinction: Prime | Active lever presses between hour 2-4 in the within-session extinction-prime reinstatement test |
| *Extinction: Day 6 | Active lever presses made during extinction session 6 |
| Extinction de-escalation | Active lever presses during extinction session 1 subtracted from session 6 |
| *Prime reinstatement | Active lever presses during prime reinstatement test |
| *Cued reinstatement | Active lever presses during cued reinstatement test |
| *Following heroin experience* |  |
| OFT distance 2 | Total distance travelled |
| OFT time 2 | Time spent moving |
| OFT difference distance | Total distance travelled during OFT 1 subtracted from OFT 2 |
| OFT time difference | Total time spent moving during OFT 1 subtracted from OFT 2 |
| EPM closed 2 | Time spent in closed arms |
| EPM open 2 | Time spent in open arms |
| EPM difference closed | Time spent in closed arm during EPM 1 subtracted from EPM 2 |
| EPM difference open | Time spent in open arm during EPM 1 subtracted from EPM 2 |
| TF baseline 2 | Average latency of tail removal during baseline (i.e., saline) session |
| TF test 2 | Average latency of tail removal during test (i.e., heroin) session |
| TF difference baseline | Average latency of tail removal during TF baseline 1 subtracted from TF baseline 2 |
| TF difference test | Average latency of tail removal during TF test 1 subtracted from TF test 2 |

**Supplemental Table 2.** Covariates for behavioral traits from rats tested at MUSC. Covariates that explained >2% of variance were regressed out prior to further analyses. Batches of 40 rats (20 males and 20 females) were shipped every few months from early 2019 through the end of 2022. Testing chamber refers to behavioral testing chamber that all heroin taking, extinction and seeking occurred in. Heterogeneous rats have 5 distinct coat colors: albino, black, brown, black hood and brown hood.

| **Trait** | **Covariates** |
| --- | --- |
| *Prior to heroin experience* |  |
| OFT distance 1 | Weight at time of surgery; Batch 6 |
| OFT time 1 | Batch 6 |
| EPM closed 1 | Weight at time of surgery; Age at time of experiment; Sex; Batch 2, 3, 4, 6, 12, 13, 15 and 17; Two specific testing chambers |
| EPM open 1 | Weight at time of surgery; Sex; Batch 2, 3, 4, 12 and 17 ; Coat color: black hood; Three specific testing chambers |
| TF baseline 1 | Batch 3 and 4; Coat color: black and brown hood |
| TF test 1 | Weight at time of surgery; Sex; Three specific testing chambers |
| *Heroin measures* |  |
| Escalation of intake: Hour 1 | None |
| Escalation of intake: Hour 1-12 | None |
| Consumption | Weight at time of surgery; Age at time of first self-administration session; Sex; Batch 14 and 15; Two specific testing chambers |
| Break point | Age at time of experiment; Behavior for Batch 14 and 15; One specific testing chamber |
| Extinction burst | Weight at time of surgery; Batch 4 and 17; 1 specific testing chamber |
| Extinction: Prime | Weight at time of surgery; Batch 2, 4, 14, 17; Two specific testing chambers |
| Extinction: Day 6 | Batch 3 |
| Extinction de-escalation | Batch 2 |
| Prime reinstatement | Weight at time of surgery; Weight at time of experiment; Sex; batch 3; One specific testing chamber |
| Cued reinstatement | Weight at time of surgery for; Age at time of experiment; Behavior for Batch 2, 3, 15 and 17; Two specific testing chambers |
| *Following heroin experience* |  |
| OFT distance 2 | Age at time of experiment for Batch 2 |
| OFT time 2 | Age at time of experiment; Batch 2 |
| OFT difference distance | Batch 6 |
| OFT time difference | Batch 6 |
| EPM closed 2 | Weight at time of surgery; Sex; Batch 2, 6, 12, 16. One specific testing chamber |
| EPM open 2 | Weight at time of surgery; Sex; Batch 2, 6, and 16; Coat color: albino and black hood; Two specific testing chambers |
| EPM difference closed | Batch 2, 3, 4, 5, 6, 7, 9, 10, 11, 13 and 16 |
| EPM difference open | Batch 7 and 16. Coat color: albino |
| TF baseline 2 | Weight at time of surgery; Batch 2, 3, 4, 8, 14 |
| TF test 2 | Weight at time of surgery; Sex; Batch 3, 4, 7; Three specific testing chambers |
| TF difference baseline | Batch 2 |
| TF difference test | Weight at time of surgery |

**Supplemental Table 3.** Covariates for behavioral traits from rats tested at UCAM. All covariates that explained >2% of variance for a trait were regressed out prior to continuing analyses. Rats were shipped every few months from early 2019 through the end of 2022 in batches of 40 rats (20 males and 20 females). Testing chamber refers to behavioral testing chamber that all heroin taking, extinction and seeking occurred in. Heterogeneous rats have 5 distinct coat colors: albino, black, brown, black hood and brown hood.

| **Trait** | **Covariates** |
| --- | --- |
| *Prior to heroin experience* |  |
| OFT distance 1 | Weight at time of surgery; Sex; Four specific testing chambers |
| OFT time 1 | Weight at time of surgery; Sex; Batch 8 and 14; Two specific testing chambers |
| EPM closed 1 | Batch 3, 9, 11, 14 and 16 |
| EPM open 1 | Weight at time of surgery; Age at time of experiment; Sex; Batch 9 and 11; One specific testing chamber |
| TF baseline 1 | Weight at time of surgery; Batch 5, 7, 8 and 14; Coat color: brown, albino and brown hood |
| TF test 1 | Weight at time of surgery; Sex; Batch 3, 4, 5, 6, 7, 13 |
| *Heroin measures* |  |
| Escalation of intake: Hour 1 | Batch 4, 5, 12 and 13 |
| Escalation of intake: Hour 1-12 | Batch 4, 5, 6, 12 and 14 |
| Consumption | Weight at time of surgery; Age at time of first self-administration session; Sex; One specific testing chambers |
| Break point | Weight at time of surgery; Sex; Batch 4 and 11 |
| Extinction burst | Weight at time of surgery; Sex |
| Extinction: Prime | Weight at time of surgery; Sex; Batch 6 |
| Extinction: Day 6 | Weight at time of surgery; Sex; Batch 6, 12 and 14; One specific testing chamber |
| Extinction de-escalation | Batch 6 |
| Prime reinstatement | Weight at time of surgery; Sex; batch 8, 10 and 12; One specific testing chamber |
| Cued reinstatement | Weight at time of surgery for; Sex; Batch 12, 13, 14; One specific testing chamber |
| *Following heroin experience* |  |
| OFT distance 2 | Weight at time of surgery; Age at time of experiment; Sex; Two specific testing chambers |
| OFT time 2 | Weight at time of surgery; Sex; Batch 8 and 16 |
| OFT difference distance | Age at time of experiment; Sex; Batch 8; Two specific testing chambers |
| OFT time difference | Age at time of experiment; Sex; Two specific testing chambers |
| EPM closed 2 | Weight at time of surgery; Batch 4 and 8 |
| EPM open 2 | Weight at time of surgery; Sex; Batch 6; One specific testing chamber |
| EPM difference closed | Batch 3 and 11 |
| EPM difference open | Two specific testing chambers |
| TF baseline 2 | Weight at time of surgery; Sex; Batch 7 and 16; Coat color: albino; One specific testing chamber |
| TF test 2 | Weight at time of surgery; Age at time of experiment; Batch 4, 5, 8, 10, 11, 12, 13, 14 and 16 |
| TF difference baseline | None |
| TF difference test | Batch 3, 5, 7, 8, 9, 10, 11, 12 and 14 |

**Supplemental Table 4.**
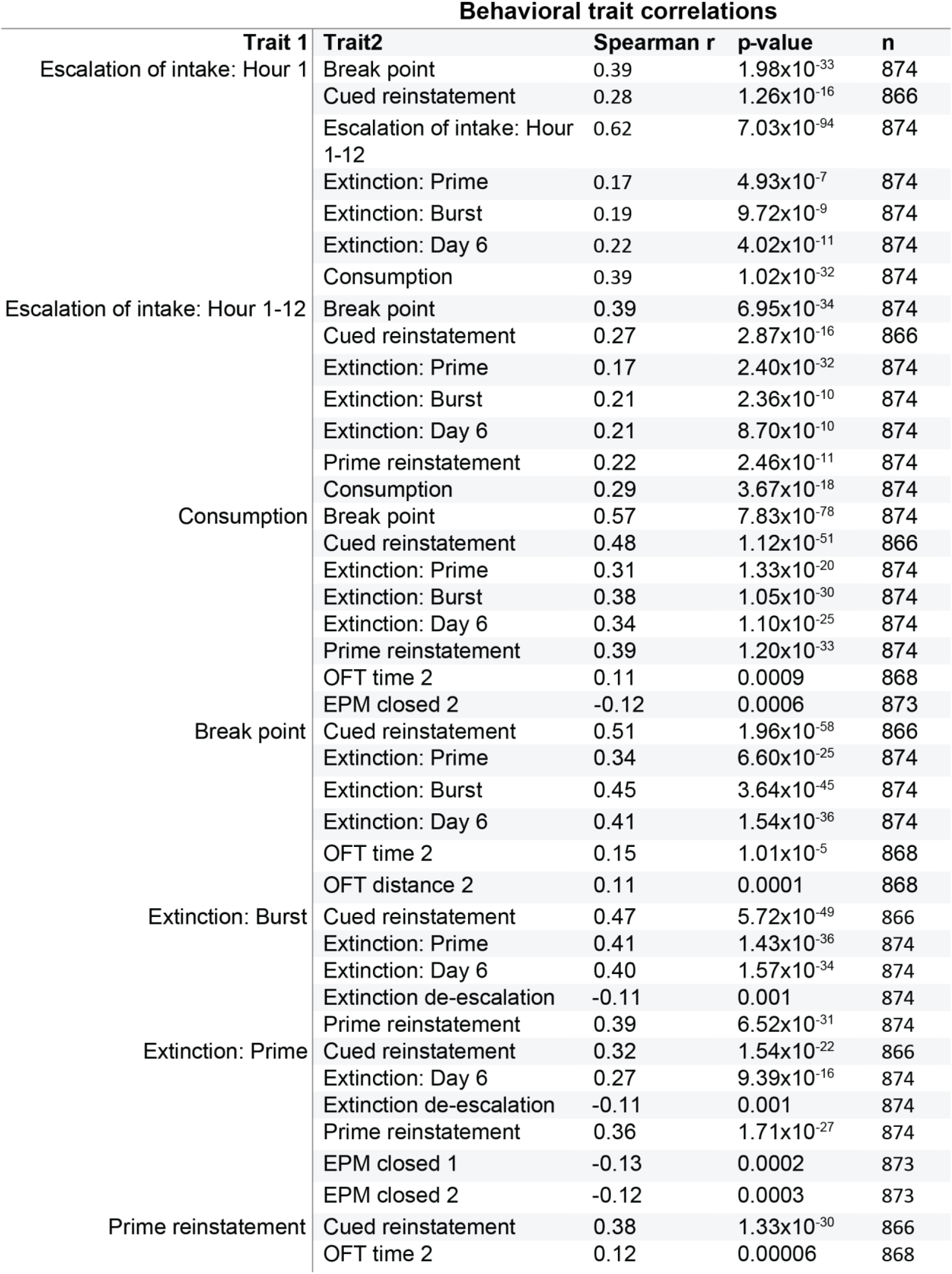
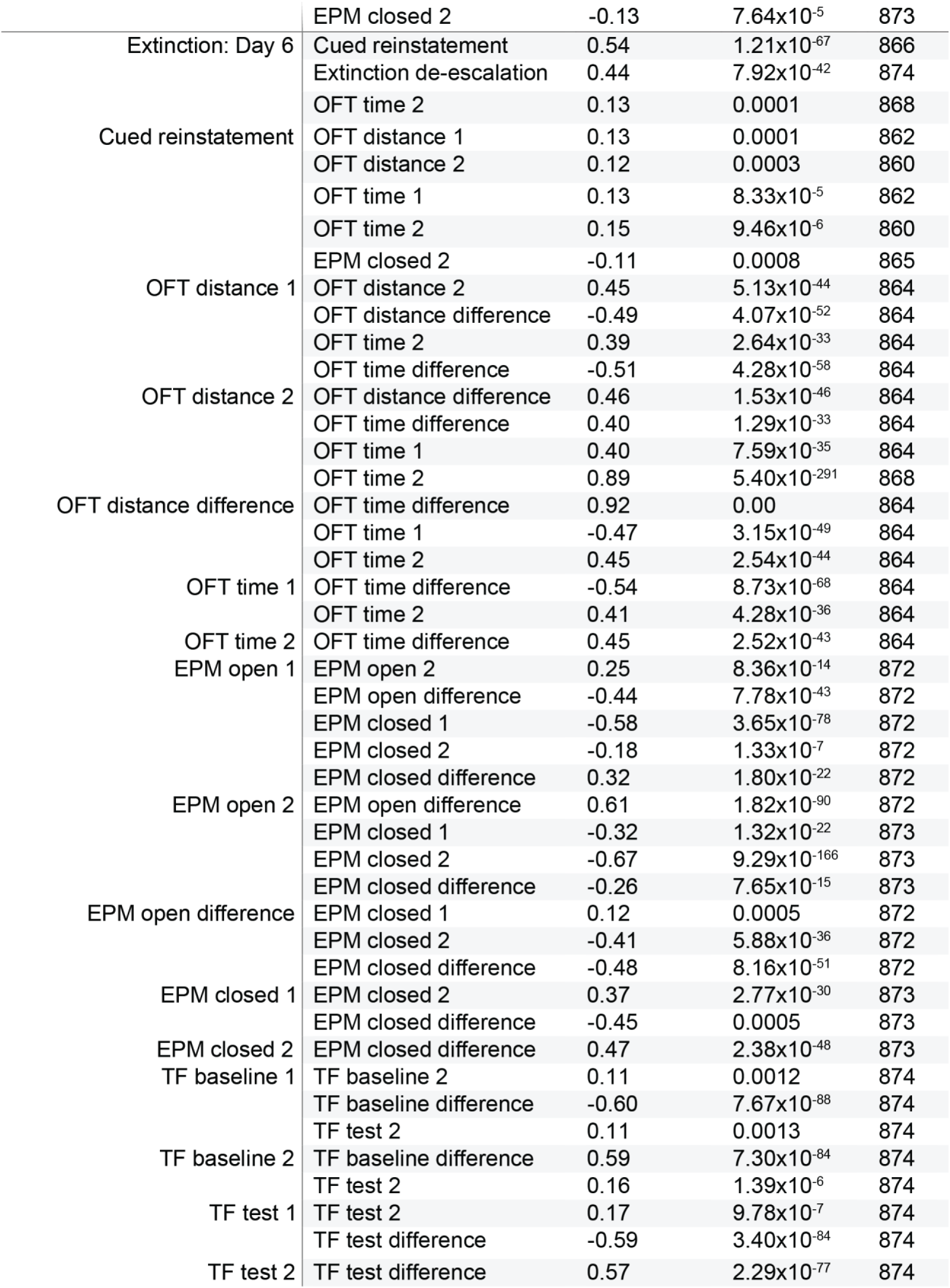
Data statistics for significant correlated behavioral traits. Heroin taking, refraining and seeking behaviors showed the tendency to co-vary with one another, whereas behaviors assessing stress- and anxiety-like behaviors as well as nociceptive threshold exhibited greater within-test covariance. Direction and magnitude of correlation (Spearman r) and total number of animals included in analysis (n) are included in the table. A significance threshold of p<0.05 with Bonferroni adjustment for correction for multiple comparisons was implemented, resulting in a p<0.0018 in order to attain significance.

**Supplemental Table 5.** SNP heritability table for behavioral traits. Significant relationships are highlighted in bold. The heritability index (h^2^SNP), standard error (SE), p-value and total number of animals in analysis (n) for each trait is recorded.

| Trait | h<sup>2</sup>SNP | SE | p-value | n |
| --- | --- | --- | --- | --- |
| OFT distance 1 | 0.139 | 0.048 | **<0.001** | 870 |
| OFT time 1 | 0.103 | 0.046 | **0.004** | 870 |
| EPM open 1 | 0.054 | 0.041 | 0.073 | 873 |
| EPM closed 1 | 0.083 | 0.043 | **0.012** | 873 |
| TF baseline 1 | 0.11 | 0.046 | **0.003** | 874 |
| TF test 1 | 0.047 | 0.039 | 0.1 | 874 |
| Escalation of intake: Hour 1 | 0.062 | 0.042 | 0.059 | 874 |
| Escalation of intake: Hour 1-12 | 0.037 | 0.04 | 0.168 | 874 |
| Consumption | 0.172 | 0.052 | **<0.001** | 874 |
| Break point | 0.009 | 0.039 | 0.42 | 874 |
| Extinction: Burst | 0.098 | 0.045 | **0.007** | 874 |
| Extinction: Prime | 0.002 | 0.036 | 0.48 | 874 |
| Extinction: Day 6 | 0 | 0.038 | 0.5 | 874 |
| Extinction de-escalation | 0.003 | 0.037 | 0.47 | 874 |
| Prime reinstatement | 0 | 0.036 | 0.5 | 874 |
| Cued reinstatement | 0.037 | 0.04 | 0.172 | 866 |
| OFT distance 2 | 0.232 | 0.052 | **<0.001** | 868 |
| OFT distance difference | 0 | 0.033 | 0.5 | 864 |
| OFT time 2 | 0.158 | 0.049 | **<0.001** | 868 |
| OFT time difference | 0.007 | 0.033 | 0.41 | 864 |
| EPM open 2 | 0.072 | 0.042 | **0.024** | 873 |
| EPM Open difference | 0 | 0.035 | 0.5 | 872 |
| EPM closed 2 | 0.016 | 0.035 | 0.31 | 873 |
| EPM closed difference | 0 | 0.034 | 0.5 | 873 |
| TF baseline 2 | 0.146 | 0.049 | **<0.001** | 874 |
| TF baseline difference | 0.044 | 0.04 | 0.12 | 974 |
| TF test 2 | 0.124 | 0.045 | **<0.001** | 874 |
| TF test difference | 0 | 0.035 | 0.5 | 874 |
| Vulnerable vs resilient/intermediate | 0.168 | 0.051 | **<0.001** | 862 |
| Intermediate vs vulnerable/resilient | 0.093 | 0.044 | **0.007** | 862 |
| Resilient vs vulnerable/intermediate | 0.15 | 0.053 | **0.002** | 862 |

**Supplemental Table 6.** Minor allele frequency and Hardy-Weinberg equilibrium values for significant QTLs. For each SNP identified, the chromosome number (CHR), missingness (MISS), genotype (count of reference/reference, reference/alternate, and alternate/alternate alleles), p-value of Hardy-Weinberg equilibrium (HWE; assumes H=0 is in HWE), minor allele frequency (MAF), non-reference allele (A1) and reference allele (A2). All SNPs passed the filters for MISS, HWE and MAF.

| SNP | CHR | MISS | Genotype | HWE | MAF | A1 | A2 |
| --- | --- | --- | --- | --- | --- | --- | --- |
| 19:8648697 | 19 | 0.00000 | 0/38/836 | 1.00000 | 0.02174 | C | A |
| 10:35103415 | 10 | 0.09725 | 44/288/457 | 0.92210 | 0.23830 | A | G |
| 2:51588876 | 2 | 0.05378 | 156/437/234 | 0.06793 | 0.45280 | C | G |
| 19:12470318 | 19 | 0.03661 | 103/371/368 | 0.54060 | 0.34260 | T | C |
| 11:35034818 | 11 | 0.01259 | 138/409/316 | 0.77620 | 0.39690 | T | A |

**Supplemental Table 7.** Comparison of GWAS results between sites. Table shows significant QTLs conserved between GWAS analysis when combining sites versus considering each side separately. Each trait, p-value (-log_10_) and chromosome location (chromosome: location) is recorded for each significant QTL. (MUSC, n=479; UCAM, n=395)

| Trait | Top SNP location | -Log<sub>(10)</sub> (p-value): MUSC | -Log<sub>(10)</sub> (p-value): UCAM | -Log<sub>(10)</sub> (p-value): Combined |
| --- | --- | --- | --- | --- |
| Escalation of intake: Hour 1-12 | 10:35103415 | 3.324 | 3.238 | 6.007 |
| Consumption | 11:35034818 | 3.559 | 3.673 | 6.493 |
| Break point | 19:8648697 | 2.918 | 3.763 | 6.298 |
| TF baseline 1 | 2:51588876 | 4.661 | 2.200 | 6.213 |
| TF baseline 1 | 19:12470318 | 2.912 | 3.646 | 5.531 |

**Supplemental Table 8.** Summarization of significant QTLs for selected traits and relevance to psychiatric disorders. The gene, effect (relationship to peak SNP), linkage disequilibrium (LD) to peak SNP, known functional role in psychiatric illnesses, model organisms used and citation for the study are recorded.

| **Total consumption** |  |  |  |  |  |
| --- | --- | --- | --- | --- | --- |
| Gene | Effect | LD with top SNP | Relevant psychiatric function | Model | Citation |
| *Brwd1* | Coding variant | 0.99 | Cigarette smoking | Human | 63-68 |
|  |  |  | Externalizing behaviors | Human | 69 |
|  |  |  | ADHD | Human | 66, 70 |
|  |  |  | Hyperthymia in bipolar disorder | Human | 71 |
| *PCP4* | Coding variant; eQTL | 0.99; 0.88 | Alcohol drinking and use disorder | Human | 73, 77 |
|  |  |  | Nicotine use withdrawal | Rodent | 78 |
|  |  |  | Cigarette smoking | Human | 63, 72 |
| *Sh3bgr* | Coding variant; eQTL | 0.99; 0.99 | Unknown |  |  |
| *ENSRNOG00000068065* | eQTL; sQTL | 0.99-1.00 | Unknown |  |  |
| *Get1* | sQTL | 0.99 | Unknown |  |  |
| **Break point** |  |  |  |  |  |
| *Phb1l2* | Coding variant | 1.0 | Alcohol use disorder (analogue) | Human | 85 |
| *Mmp15* | eQTL | 1.0 | MMP2 necessary for heroin extinction | Rodent | 88 |
| **Baseline nociception prior to heroin experience (Chromosome 2)** |  |  |  |  |  |
| *Ccdc152* | Coding variant | 0.77 | Unknown |  |  |
| *Ghr* | Coding variant | 0.77 | Unknown |  |  |
| *Zfp131* | Coding variant | 0.82 | Unknown |  |  |
| *Selenop* | eQTL; sQTL | 0.66; 0.78-79 | Methamphetamine toxicity | Rodent | 91 |
| **Baseline nociception prior to heroin experience (Chromosome 19)** |  |  |  |  |  |
| *Large1* | Coding variant; eQTL | 0.75; 0.69-0.75 | Educational attainment | Human | 89-91 |
|  |  |  | Cigarette smoking | Human | 63 |
|  |  |  | Alcohol dependence | Human | 92 |
|  |  |  | Neuroticism | Human | 93 |
|  |  |  | Depression | Human | 93-94 |
| *Rps18l2* | eQTL | 0.98 | Unknown |  |  |
| *Tom1* | sQTL | 1.0 | Cigarette smoking | Human | 72 |
| *Mt3* | sQTL | 0.75 | Alzheimer's and Huntington's disease | Human | 85 |

**Supplemental Table 9.**
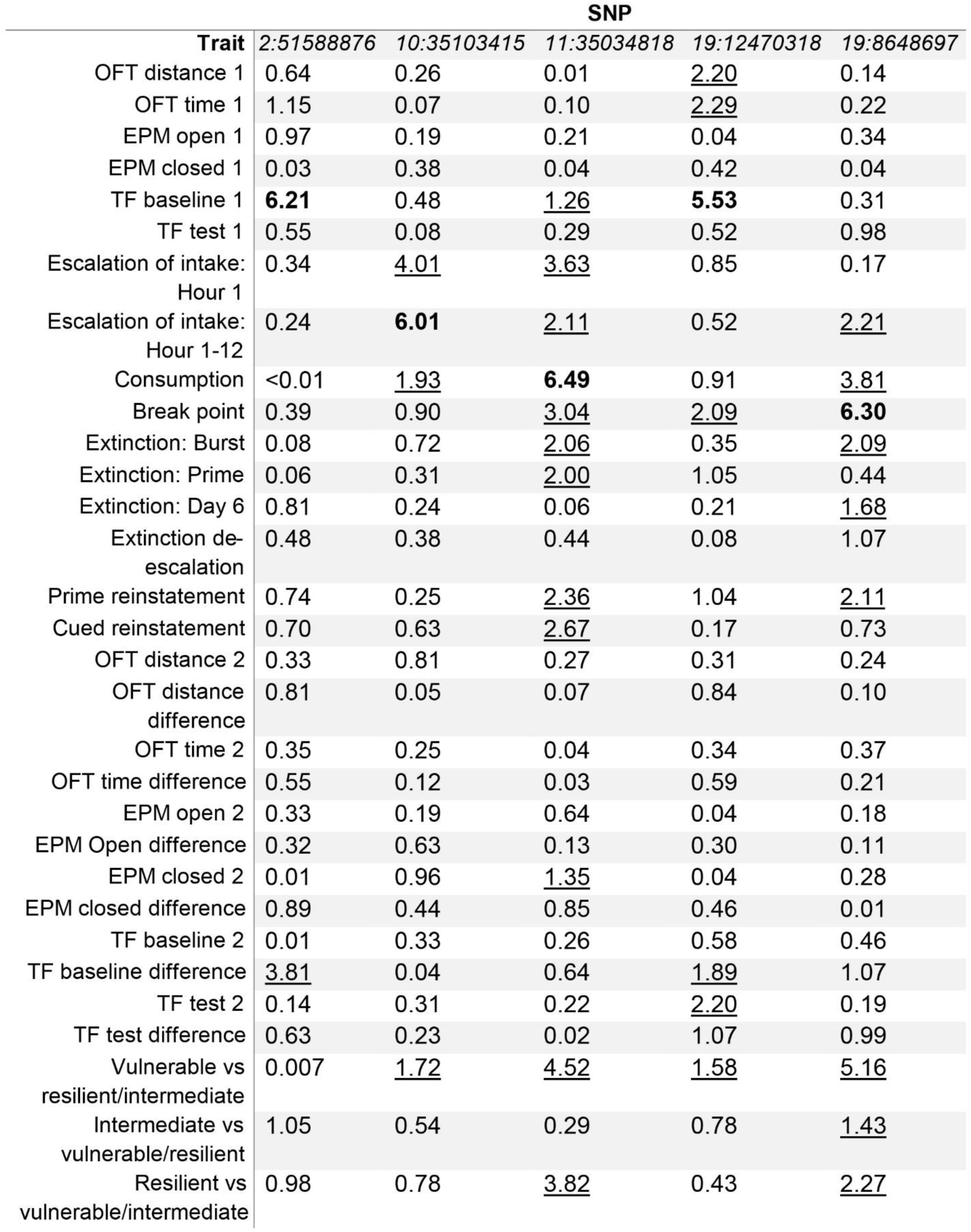
Significant SNPs across all behavioral traits tested. The p-value (-log_10_) for the significant SNPs identified in the GWAS analysis across all behavioral traits. Values in bold indicate genome-wide significant p-values; underlines indicate p-values greater than −log(p) = 1.3 (-log(p) of 1.3 = 0.05) but below the threshold for genome-wide significance of −log(p)=5.58 (-log(p) = 5.58 = 0.00000263)

|  | SNP |  |  |  |  |
| --- | --- | --- | --- | --- | --- |
| Trait | 2:51588876 | 10:35103415 | 11:35034818 | 19:12470318 | 19:8648697 |
| OFT distance 1 | 0.64 | 0.26 | 0.01 | <u>2.20</u> | 0.14 |
| OFT time 1 | 1.15 | 0.07 | 0.10 | <u>2.29</u> | 0.22 |
| EPM open 1 | 0.97 | 0.19 | 0.21 | 0.04 | 0.34 |
| EPM closed 1 | 0.03 | 0.38 | 0.04 | 0.42 | 0.04 |
| TF baseline 1 | **6.21** | 0.48 | <u>1.26</u> | **5.53** | 0.31 |
| TF test 1 | 0.55 | 0.08 | 0.29 | 0.52 | 0.98 |
| Escalation of intake:<br>Hour 1 | 0.34 | <u>4.01</u> | <u>3.63</u> | 0.85 | 0.17 |
| Escalation of intake:<br>Hour 1-12 | 0.24 | **6.01** | <u>2.11</u> | 0.52 | <u>2.21</u> |
| Consumption | <0.01 | <u>1.93</u> | **6.49** | 0.91 | <u>3.81</u> |
| Break point | 0.39 | 0.90 | <u>3.04</u> | <u>2.09</u> | **6.30** |
| Extinction: Burst | 0.08 | 0.72 | <u>2.06</u> | 0.35 | <u>2.09</u> |
| Extinction: Prime | 0.06 | 0.31 | <u>2.00</u> | 1.05 | 0.44 |
| Extinction: Day 6 | 0.81 | 0.24 | 0.06 | 0.21 | <u>1.68</u> |
| Extinction de-escalation | 0.48 | 0.38 | 0.44 | 0.08 | 1.07 |
| Prime reinstatement | 0.74 | 0.25 | <u>2.36</u> | 1.04 | <u>2.11</u> |
| Cued reinstatement | 0.70 | 0.63 | <u>2.67</u> | 0.17 | 0.73 |
| OFT distance 2 | 0.33 | 0.81 | 0.27 | 0.31 | 0.24 |
| OFT distance<br>difference | 0.81 | 0.05 | 0.07 | 0.84 | 0.10 |
| OFT time 2 | 0.35 | 0.25 | 0.04 | 0.34 | 0.37 |
| OFT time difference | 0.55 | 0.12 | 0.03 | 0.59 | 0.21 |
| EPM open 2 | 0.33 | 0.19 | 0.64 | 0.04 | 0.18 |
| EPM Open difference | 0.32 | 0.63 | 0.13 | 0.30 | 0.11 |
| EPM closed 2 | 0.01 | 0.96 | <u>1.35</u> | 0.04 | 0.28 |
| EPM closed difference | 0.89 | 0.44 | 0.85 | 0.46 | 0.01 |
| TF baseline 2 | 0.01 | 0.33 | 0.26 | 0.58 | 0.46 |
| TF baseline difference | <u>3.81</u> | 0.04 | 0.64 | <u>1.89</u> | 1.07 |
| TF test 2 | 0.14 | 0.31 | 0.22 | <u>2.20</u> | 0.19 |
| TF test difference | 0.63 | 0.23 | 0.02 | 1.07 | 0.99 |
| Vulnerable vs<br>resilient/intermediate | 0.007 | <u>1.72</u> | <u>4.52</u> | <u>1.58</u> | <u>5.16</u> |
| Intermediate vs<br>vulnerable/resilient | 1.05 | 0.54 | 0.29 | 0.78 | <u>1.43</u> |
| Resilient vs<br>vulnerable/intermediate | 0.98 | 0.78 | <u>3.82</u> | 0.43 | <u>2.27</u> |

**Supplemental Figure 1.**
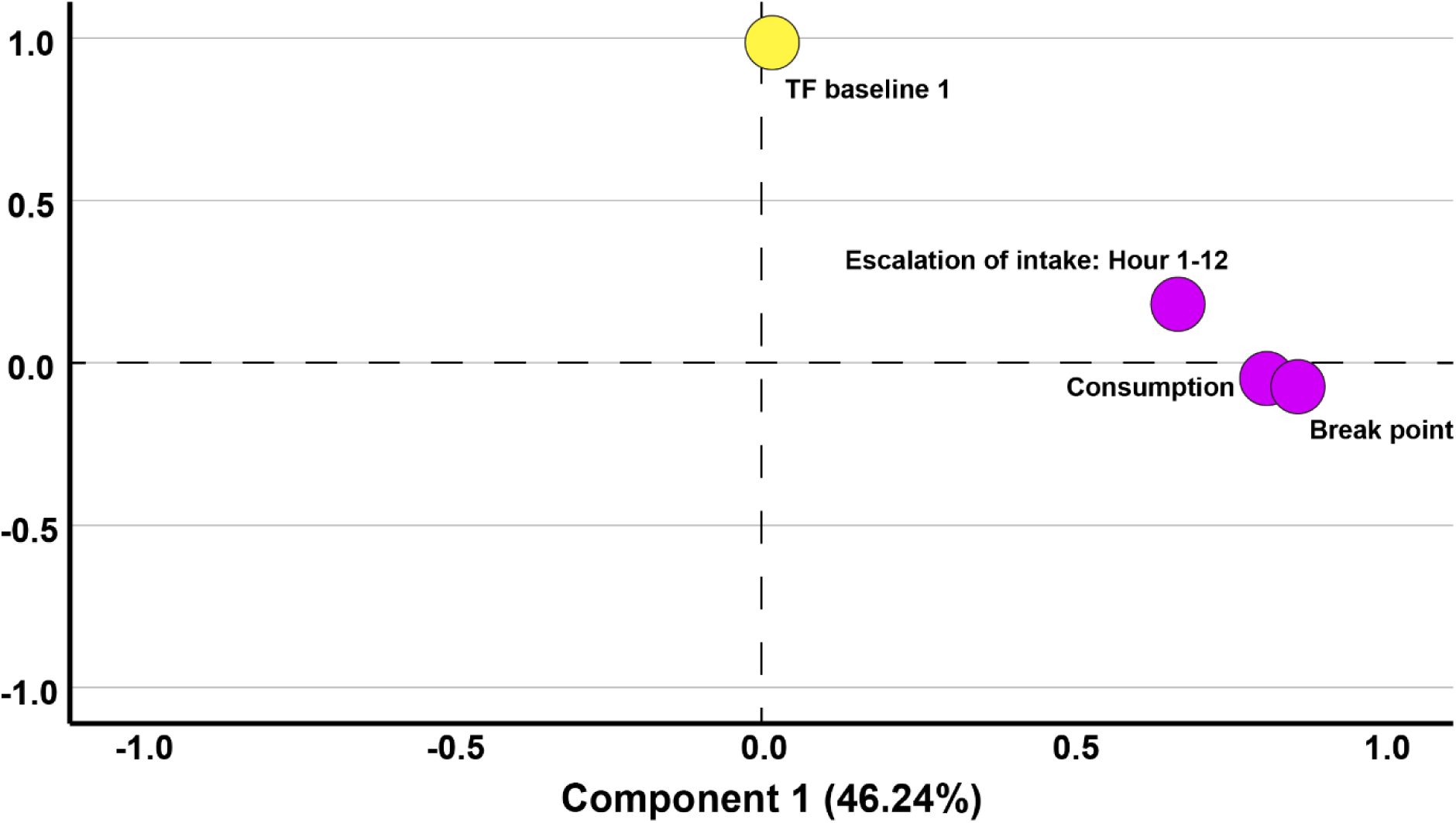
Principal component analysis for behavioral traits with significant QTLs. Two factors were identified and comprised 71.55% of total variance in the data. Heroin-taking behaviors accounted for 46.24% of variance (component 1, purple) with traits exhibiting strong loading (Break point: 0.86; Consumption: 0.81; and Escalation of intake: Hour 1-12: 0.67), and was mutually exclusive from baseline nociception prior to heroin experience (component 2, yellow) which accounted for 25.31% of variance with strong loading (0.99). Axes represent factor loading values for each trait within respective component. (n=874)

**Supplemental Figure 2.**
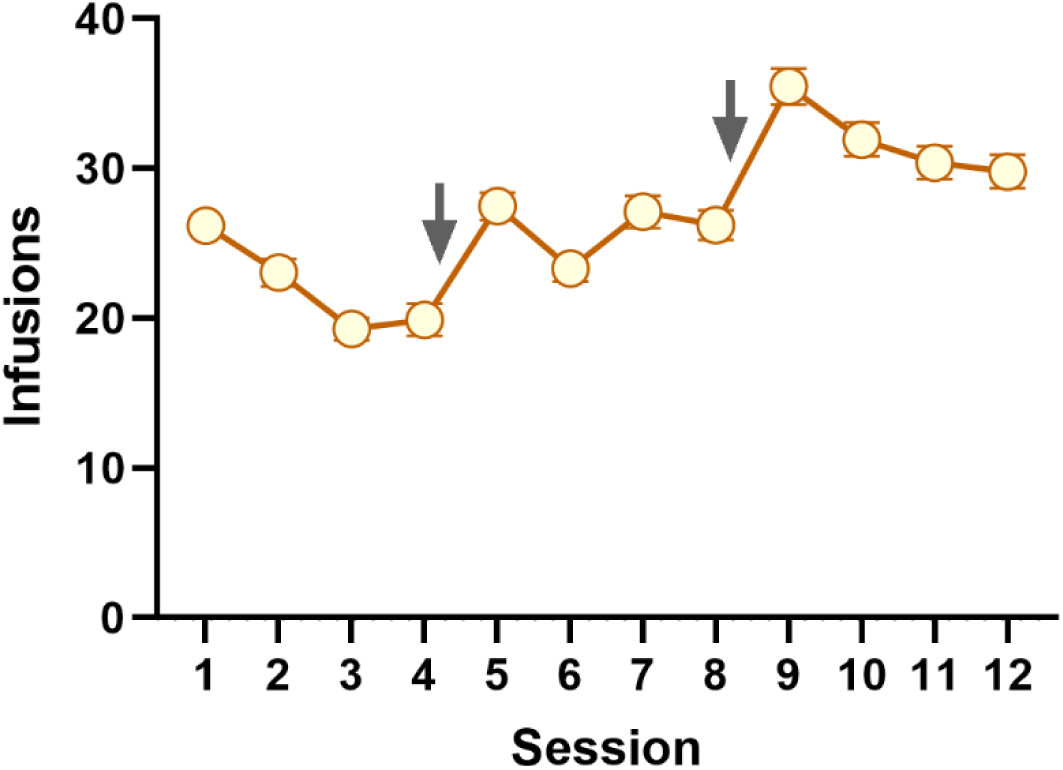
Total number of heroin infusions earned across training sessions. Mean ± SEM for total heroin infusions (20 µg/kg/100 µl infusion) earned during heroin self-administration across training sessions. Animals underwent four training sessions a week with one day random day off between Monday-Friday. Grey arrows indicate brief periods of forced abstinence between training weeks. Rats increased heroin intake over the course of training (mixed effects ANOVA; F(5.90,5399)=43.24, p<0.001). (n=874)

